# mTORC3 activity is regulated by GSK3β through the activation of PKCδ that phosphorylates and activates STAT1

**DOI:** 10.1101/2024.01.26.577410

**Authors:** Jun Zhan, Frank C. Harwood, Monica Cardone, Gerard C. Grosveld

## Abstract

The mechanistic target of rapamycin serine/threonine kinase (mTOR), a critical regulator of cell proliferation, is implicated in many human diseases, including cancer. Our laboratory recently discovered a new mTOR complex, called mTORC3, distinct from the two canonical mTOR complexes, mTORC1, and mTORC2, which causes accelerated proliferation and increased malignancy of cancer cells. Understanding how mTORC3 is controlled is an important approach to designing cancer therapies targeting mTORC3. Having investigated how mTORC3 activity was affected by glycogen synthase kinase-3β (GSK3β), a known regulator of mTOR activity, we found that cells do not assemble mTORC3 upon GSK3β knockout, due to extinguished expression of ETV7, an essential component of the mTORC3 complex. We discovered that the lack of ETV7 expression resulted from reduced phosphorylation of serine 727 (S727) of the Signal Transducer and Activator of Transcription 1 (STAT1), which is required for transcriptional activation of ETV7. Finally, we identified that GSK3β through PKCδ phosphorylates STAT1-S727. Together, we identified a novel GSK3β/PKCδ/ STAT1 pathway in Karpas cells that regulates ETV7 transcription, thus positively regulating mTORC3 activity. The identification of this signaling pathway could benefit the therapeutic design for numerous diseases including cancer.

## Introduction

mTOR is a serine/threonine kinase that regulates both cell survival and cell growth through its ability to integrate signals from nutrients and growth factors[1–3]. mTOR signaling functions through two structurally and functionally distinct complexes, mTOR complex 1 (mTORC1) and mTOR complex 2 (mTORC2), which are activated as a consequence of ligand activation of growth factor receptors[4]. mTORC1 phosphorylates the protein synthesis regulators P70S6K and 4E-BP1 to control protein translation[2], while mTORC2 phosphorylates members of the AGC family of kinases such as AKT, members of the protein kinase C family (PKCs)[2], and serum- and glucocorticoid-induced protein kinase-1 (SGK-1)[2]. mTORC2 phosphorylation of AGC kinases induces the activation of multiple downstream targets, which in turn affect cell fate and function[5]. Recently, we have identified a new mTOR complex that is distinct from the mTORC1 and mTORC2 complexes, called mTORC3[6], which phosphorylates substrates of both mTORC1 and mTORC2. An essential component of mTORC3 is the ETS transcription factor ETV7[7]. Cells that assemble ETV7/mTORC3 grow fast and acquire resistance to rapamycin. Disruption of mTORC3 assembly might be an important strategy for cancer therapy.

Another important and extensively investigated therapeutic target for cancer is glycogen synthase kinase-3β (GSK3β), which has been reported to crosstalk with mTOR signal transduction pathways. GSK3β is a serine/threonine kinase that was initially identified as a key protein in glucose metabolism[8, 9]. However, GSK3β regulates many cellular functions, including cell proliferation, cell survival, gene expression, cellular architecture, neural development, and cell plasticity[10, 11]. To reflect its multiple functions, GSK3β phosphorylates many different protein substrates that are components of multiple key cellular pathways, including the Wnt/β-catenin, Hedgehog, Notch, NF-κB, and PI3K/Akt[12–15] pathways. Both GSK3β overexpression and high GSK3β activity have been found in many cancers; therefore, it is a positive regulator of cancer cell proliferation and survival[16–18]. GSK3β accomplishes this by positively regulating mTOR activity through phosphorylation of the mTOR-associated scaffold protein Raptor (regulatory-associated protein of mTOR) on S859[19], and through phosphorylation and subsequent degradation of REDD1, an inhibitor of mTOR[20]. To counterbalance this effect, GSK3β activity is negatively regulated by mTOR through AKT[21] or p70S6K[22].

In this paper, we show that GSK3β indirectly regulates ETV7 expression thereby influencing mTORC3 activity. We show that ETV7 transcription is activated by STAT1, whose activity is regulated by GSK3β through phosphorylation of STAT1-S727 by PKCδ.

## Materials and methods

### Antibodies

Anti-phospho-P70S6K-T389, anti-P70S6K, anti-STAT1, anti-phospho-STAT1-S727, anti-phospho-STAT1-Y701, anti-GSK3β, anti-phospho-GSK3β-Y216, anti-PKCδ, anti-PKCδ-Y311, anti-phospho-β-catenin-S33/S37/T41, anti-β-catenin, anti-ERK1/2, anti-phospho-ERK-T202/Y204, anti-STAT1 (Cut & Run certified #14994) were purchased from Cell Signaling Technologies (Danvers, MA); anti-SGK1, anti-phospho-SGK-S422, were from Thermo Fisher Scientific (Waltham, MA). The ETV7 rabbit polyclonal was obtained from Sigma-Aldrich (Saint Louis, MO).

### Plasmids

For GSK3β disruption, a GSK3β guide RNA (GTCCTGCAATACTTT<u>CTT</u>GA, the underlined CTT codon encodes K85 on the coding strand) and scaffold was purchased from IDT (Integrated DNA Technologies, Coralville, IA) and cloned in LentiCRISPR-V2. For cloning of the GSK3β lentiviral vector, GSK3β cDNA was synthesized from IDT and inserted into pCL20-EF1p-ETV7-T2A-Blast using the BsiWI/BsrGI sites, yielding pCL20-EF1p-GSK3β-T2A-Blast. For cloning of the STAT1 genomic editing vector, double-stranded DNA encoding the STAT1 guide RNA (TCCCATTACAGGCTCAGTCG, targeting exon 6) and scaffold was purchased from IDT and inserted into Lenti-CRISPR-V2-noCAS using the KpnI/EcoRI sites, yielding Lenti-CRISPR-V2 STAT1sgRNA.

### Generation of Karpas-299 GSK3β and STAT1 gene knockout clones

To create gene deletions in Karpas-299 cells, we have used a CAS9-expressing cell line, called Karpas-299–EBNA–Cas9[6]. Karpas-299–EBNA–Cas9 cells (1 × 10^6^ cells) were transduced with a lentiviral-vector encoding guide RNAs for GSK3β or STAT1. Twenty-four hours later, cells were selected for 5 days with blasticidin (1 μg/ml) and then seeded (1000 cells per 10-cm dish) in methylcellulose [1.5% methylcellulose, RPMI-1640, 10% FBS, uridine (7 μg/ml), and thymidine (40 ng/ml)]. Two weeks later, 20 clones were selected, expanded in liquid culture, and evaluated by Western blotting or DNA sequencing to confirm gene knockout.

### Cell culture

Karpas-299, Karpas-299-EBNA-Cas9, and Karpas-299-K85 cells (C1, C2, and C3) were maintained in RPMI-1640 medium (Corning), supplemented with 10% bovine calf serum (BCS) (Thermo Fisher Scientific), 50 mM GlutaMAX (Life Technologies), penicillin (100 U/ml), and streptomycin (100 µg/ml) (Life Technologies) in a humidified incubator at 37°C and 5% CO_2_.

### Western blotting

Cells for Western blot analysis were lysed in 1× Cell Signaling lysis buffer [20 mM tris-HCl, (pH 7.5), 150 mM NaCl, 1 mM Na2EDTA, 1 mM EGTA, 1% Triton X-100, 2.5 mM sodium pyrophosphate,1 mM β-glycerophosphate, 1 mM Na3VO4, and leupeptin (1 µg/ml) (Cell Signaling Technologies, Thermo Fisher)] supplemented with 1 mM phenylmethylsulfonyl fluoride (PMSF). The lysates were loaded on pre-cast 4 to 12% bis-tris protein gels (Life Technologies). Proteins were transferred to nitrocellulose membranes using the iBLOT system (Life Technologies) following the manufacturer’s protocol. Membranes were blocked with 5% milk and 0.1% Tween-20 in tris-buffered saline (TBS) and incubated with the appropriate antibodies in 5% bovine serum albumin in TBS with 0.1% Tween 20, overnight at 4°C. All primary antibody incubations were followed by incubation with secondary horseradish peroxidase (HRP)– conjugated antibody (Pierce, Thermo Fisher) in 5% milk and 0.1% Tween 20 in TBS and visualized using Super Signal West Pico or Femto Chemiluminescent Substrate (Pierce, Thermo Fisher) on a ChemiDoc-MP imaging system (Bio-Rad).

### Kinase assays

Kinase assays were performed in kinase buffer (25 mM HEPES, pH 7.4, and 50 mM KCl). Just before use we added, 10 mM MgCl_2_, 10 nM of the PP1/2 inhibitor calyculin A, 0.5 mM DTT, and 1 μM of the MEK1/2 inhibitor UO126. Purified recombinant STAT1 protein was used as a substrate (Abcam, Cambridge, UK). Prior to performing the kinase assay, STAT1 proteins were dephosphorylated by lambda phosphatase (NEB, Ipswich MA) followed by the addition of protease and phosphatase inhibitors (Thermo Fisher). Purified active mTOR kinase domain(MiliporeSigma, Burlinton, MA), PKCδ (R & D Systems, Minneapolis, MN), or GSK3β recombinant protein (Abcam, Cambridge, UK) was added to the dephosphorylated protein substrates in the presence of 250 μM ATP and incubated for 30 min at 30°C. Kinase reactions were terminated by adding 4X LDS loading buffer (NuPAGE LDS, Invitrogen, Carlsbad, CA) and loaded on precast 4 to 12% bis-tris gels (Life Technologies/Thermo Fisher). The kinase substrates were visualized using a phospho-specific antibody against pSTAT1-S727.

### mRNA extraction and qRT-PCR

RNA was extracted from 0.5 - 1 million cells with the RNeasy kit (Invitrogen). One-step q-RT-PCR was performed. The ETV7 mRNAs were quantified by Taqman qPCR, with the amount of ETV7 mRNAs normalized against the amount of GAPDH mRNA. Quantitative RT-PCR was performed with TaqMan fast universal PCR master mix (Applied Biosystems; Foster City, CA). RT reaction conditions: 48°C for 30min, 95°C for 10min. PCR reaction conditions: 40 cycles of 1 second 95°C denaturation, 20 sec 60°C annealing/elongation, followed by a 5 min 72°C hold.

### STAT1 Cut and Run/PCR analysis of the ETV7 promoter

To show that STAT1 binds to the ETV7 promoter in Karpas-299 cells, we performed a Cut and Run/Q-PCR (C&R/Q-PCR) experiment of 5×10^5^ untreated cells and 5×10^5^ cells treated for 30 minutes with recombinant human INFγ (STEMCELL Technologies, Vancouver, BC, Canada). For the C&R procedure, we used the kit of EpiCypher and an anti-STAT1 antibody (#14994, Cell Signaling) or a non-specific IgG as a control, following the methods described in the manufacturer’s protocol. The amount of ETV7 promoter fragment in the IgG and anti-STAT1 C&R was determined using the Q-PCR kit of Cell Signaling following the manufacturer’s protocol. Q-PCR was performed using a BioRad CFX Opus 96 Q-PCR machine (Hercules, CA). Sequences of the Q-PCR primers: Forward primer -TCGTTTCCCGGAAGGATAG-, Reverse primer - GCGTGGGAGGAATCTCTTT-. The detection of Q-PCR products (110 bp) was performed using BioRad’s SYBR Green Supermix.

## Results

### Disruption of GSK3β leads to poor cell growth and abolishes ETV7 expression

Like mTORC1, GSK3β can directly phosphorylate the 70 kDa ribosomal protein S6 Kinase 1 (p70S6K1) [23], and 4EBP1[24], and activate the downstream signaling cascade of mTORC1 to enhance protein synthesis and cell proliferation[25]. To investigate how GSK3β affects mTORC3 activity, we disrupted the GSK3β gene in Karpas-299 cells using CRISPR/Cas9. We focused on lysine 85 of GSK3β which is essential for its kinase activity[26]. Initially, one clone with GSK3β disruption was established, called K85-C1 (Fig. 1a top lanes). Genomic sequencing covering the disrupted region of GSK3β confirmed the elimination of active GSK3β in both alleles (Supplemental Fig. 1a, 1b). Subsequently, K85-C1 cell proliferation was measured. Compared to its Karpas-299 parental cell line, K85-C1 cells grew considerably slower with a doubling time of almost 48 hours versus less than 24 hours for that of Karpas-299 (supplemental Fig. 1c). Flow cytometric analysis showed a reduced percentage of K85-C1 cells in S-phase (36%) compared to that in its parent cell line Karpas -299 (53%) (Fig. 1b).

**Figure 1.**
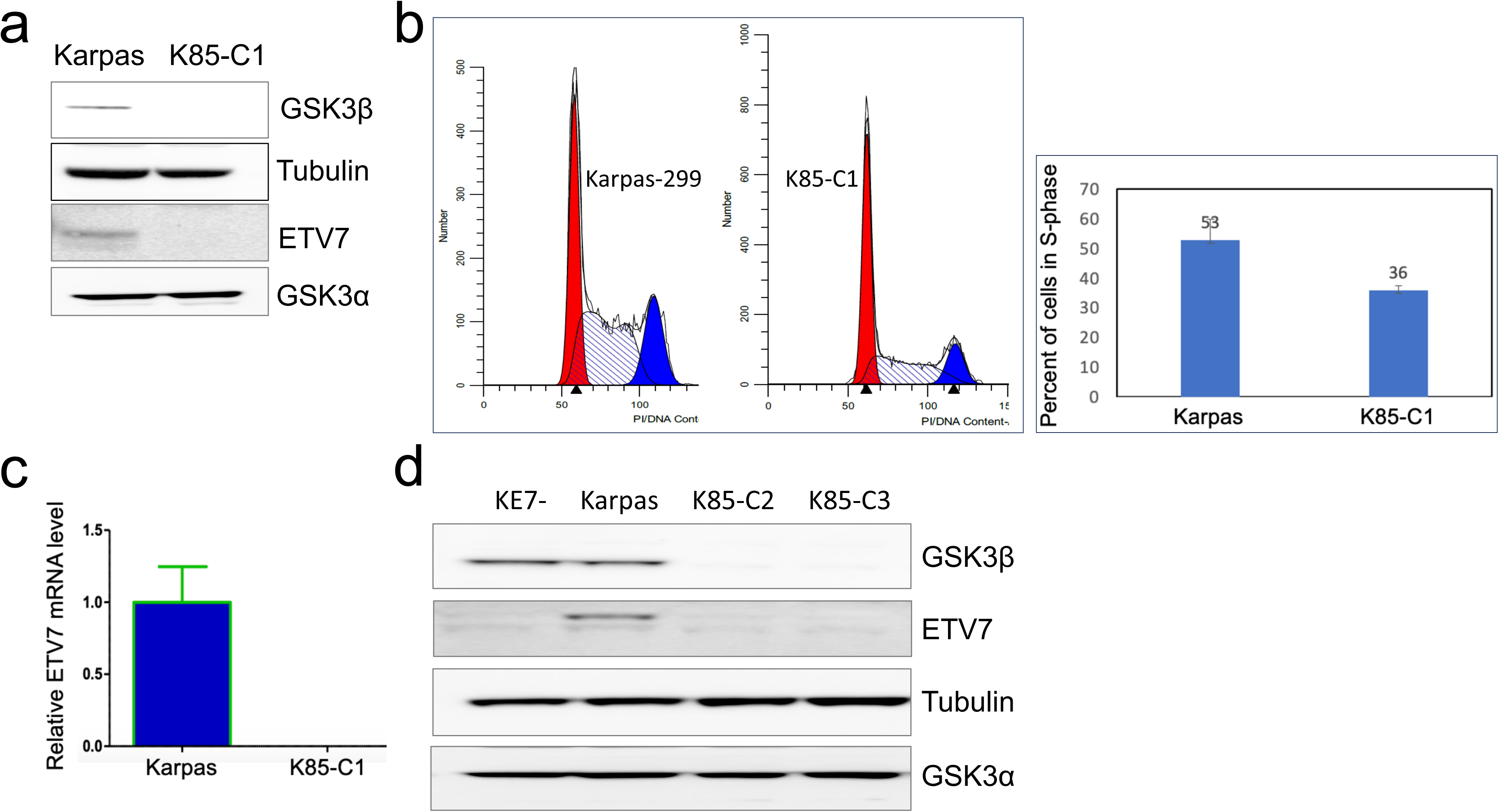
Disruption of GSK3β leads to poor cell growth and extinction of ETV7 expression. (**a**) Immunoblot of lysates from equal numbers of Karpas-299 and K85-C1 cells probed for GSK3β, tubulin, ETV7 and GSK3α. (**b**) Flow cytometric cell cycle analysis of Karpas-299 and K85-C1 cells. The left panel represents the flow chard, and the right panel shows the percentage of cells in S-phase with standard deviation between three independent experiments. (**c**) Relative amounts of ETV7 mRNA in Karpas-299 and K85-C1 cells were determined by real-time RT-PCR. (**d**) Immunoblot of lysates from equal numbers of Ke7-, Karpas-299 and clones K85-C2 and K85-C3 probed for GSK3β, tubulin, ETV7 and GSK3α.

As mTORC3 accelerates cell proliferation, we determined whether the K85-C1 cells had lost mTORC3 kinase activity. To this end, we investigated the expression of ETV7, an essential component for mTORC3 assembly. Indeed, the expression of ETV7 in K85-C1 was completely abolished (Fig. 1a), with loss of mTORC3, suggesting that GSK3β is directly or indirectly essential for the activity of mTORC3. Next, we investigated whether GSK3β affects ETV7 expression in K85-C1 cells at the transcriptional (RNA) or translational (protein) level. Quantitative real-time RT-PCR of whole cell RNA from Karpas-299 and K85-C1 cells showed that there was no ETV7 mRNA in K85-C1 cells (Fig. 1c), suggesting that GSK3β affects ETV7 expression transcriptionally.

To further validate the link between the loss of GSK3β and ETV7 expression in Karpas-299 cells, we have screened and identified additional GSK3β knockout clones, i.e. K85-C2 and K85-C3. In these clones there is neither ETV7 protein expression (Fig 1d), nor is there any ETV7 mRNA (Supplemental Fig. 1d), thus strengthening the connection between GSK3β and ETV7 expression and mTORC3 activity in Karpas-299 cells. Consistent with the above, the additional K85 clones grow slowly like K85-C1, with a reduced percentage of cells in S-phase (Supplemental Fig. 1e).

### The transcriptional activity of STAT1 is essential for the expression of ETV7

We hypothesized that GSK3β kinase activity likely phosphorylates one or more transcription factors, which, in turn, regulate ETV7 expression. A search of the Eukaryotic Promoter Database (EPD, https://epd.epfl.ch//index.php) showed that the *ETV7* gene has two promoters 116 bp apart, ETV7-P1 (proximal) and ETV7-P2 (distal), of which P2 contains a high probability (p = 0.00001) STAT1 binding site, as scored by the JASPAR CORE 2018 transcription factor binding site motif database (http://jaspar.genereg.net/) (Supplemental Fig. 2a). This implies a potential role of STAT1 in ETV7 expression. Next, we determined whether GSK3β deletion affected the STAT1 protein level and/or its phosphorylation status. Although we found that the level of STAT1 protein remained unchanged in K85-C1 compared to Karpas-299 cells (Fig. 2a), phosphorylation of STAT1-S727 was severely reduced, while the STAT1-Y701 phosphorylation was also somewhat reduced (Fig. 2a). These data were confirmed in the additional K85 cell clones (Fig. 2b). Because phosphorylation of STAT1-Y701 is required for STAT1’s nuclear translocation and phosphorylation of STAT1-S727 for its full transcriptional activation[27, 28], loss of the latter would severely reduce STAT1’s transcriptional activity in K85 cells.

**Figure 2.**
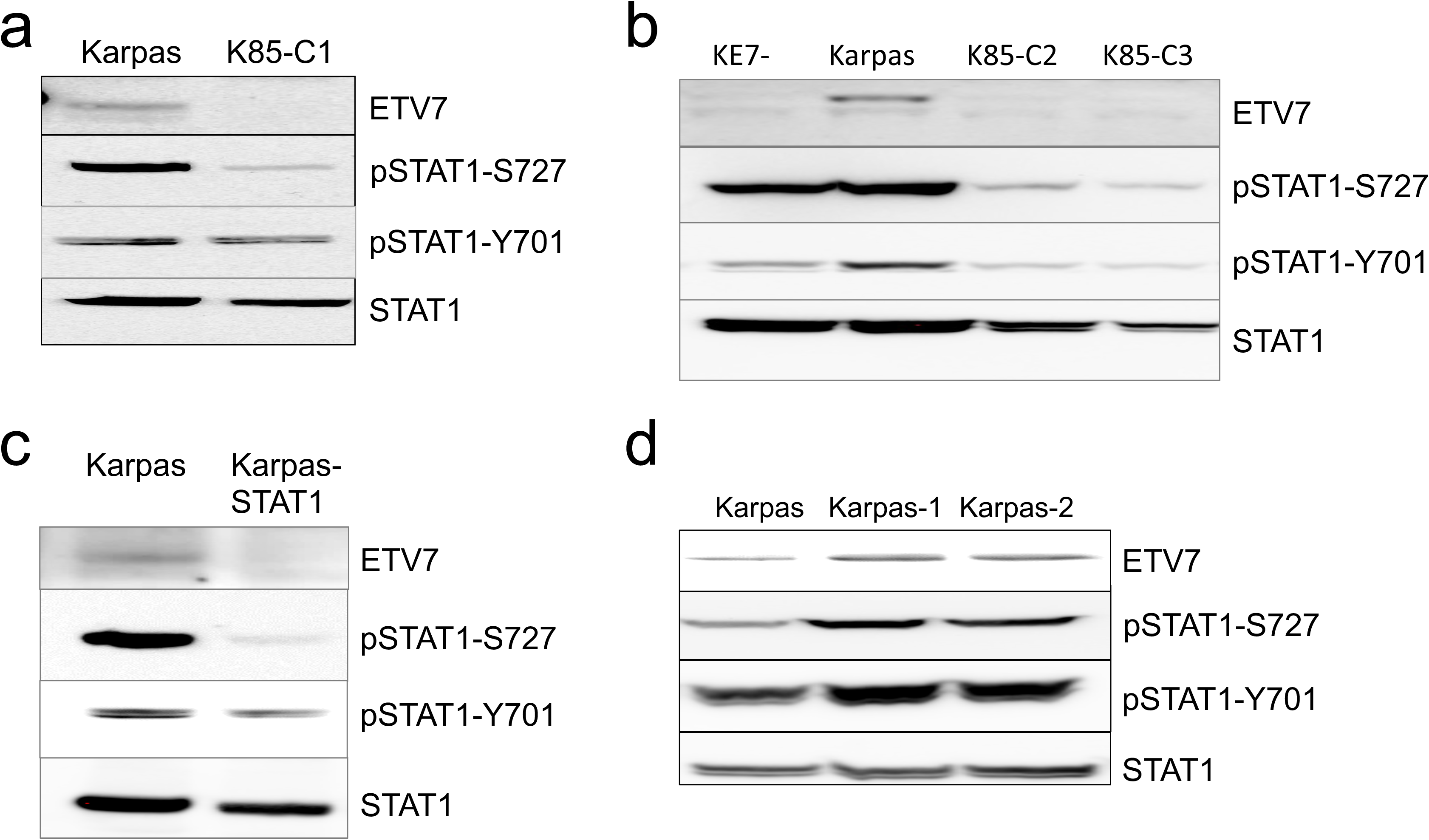
STAT1 transcriptional activity is essential for ETV7 expression. (**a**) Immunoblots of lysates from equal numbers of Karpas-299 and K85-C1 cells probed for ETV7, STAT1, pSTAT1-S727, and pSTAT1-Y701. (**b**) Immunoblots of lysates from equal numbers of KE7-, Karpas-299 and K85-C2 and K85-C3 probed for ETV7, STAT1, pSTAT1-S727, and pSTAT1-Y701. (**c**) Immunoblots of lysates from equal numbers of Karpas-299 and Karpas-299 STAT1-edited cells probed for ETV7, STAT1, pSTAT1-S727, pSTAT1-Y701. (**d**) Immunoblots of lysates from equal numbers of Karpas-299 and different increased ETV7 expressing Karpas-299 cell pools (Karpas-1, Karpas-2) probed for ETV7, STAT1, pSTAT1-S727, pSTAT1-Y701.

To further validate that STAT1 regulates the expression of ETV7 in Karpas-299 cells, we gene-edited *STAT1* in Karpas-299 cells using CRISPR-Cas9, to delete the *STAT1* gene. We failed to obtain any *STAT1* knockout or mutated cell clones as indicated by genomic sequencing of the targeted region and sequencing of the whole cDNA.

Surprisingly, we obtained clones that showed severely reduced levels of STAT1-S727 phosphorylation for unknown reasons. Using these clones, we again observed decreased ETV7 protein expression (Fig. 2c) as well as reduced ETV7 mRNA expression (Supplemental Fig. 2b), suggesting that STAT1 is a transcription factor required for *ETV7* transcription. The role of STAT1 in *ETV7* regulation was further confirmed by the selection of two single-picked Karpas-299 cell clones, that showed increased STAT1 activity, as indicated by the increased level of phosphorylation at S727 and Y701 of STAT1. Compared with the Karpas-299 cell line, these clones (Karpas-1 and Karpas-2) showed an increased level of ETV7 expression both at the protein (Fig. 2d) and the mRNA level (Supplemental Fig. 2c). Furthermore, STAT1 ChIP-seq analysis of K562 cells in the ENCODE database showed that STAT1 binds to the ETV7 P2 promoter (Supplemental Fig. 2d). A Cut and Run/Q-PCR analysis of Karpas-299 cells treated with INFγ or not, using a STAT1 antibody, showed binding of STAT1 to the ETV7 promoter in this cell line and increased STAT1 binding upon INFγ treatment (Supplemental Fig. 2e). Lastly, the conserved role of the STAT1 site in the P2 promoter is further supported by its full sequence preservation among different classes of vertebrates (Supplemental Fig. 2f). Together our data indicate that activated STAT1 is essential for ETV7 expression.

### mTOR is not responsible for STAT1-S727 phosphorylation

Next, we investigated how GSK3β regulates STAT1-S727 phosphorylation. In K85 cells, we observed reduced mTORC1 activity, as indicated by the level of phosphorylation of p70S6K-T389 (Fig. 3a), indicating that, as reported, GSK3β positively regulates mTOR activity. This suggested that mTOR, in addition to GSK3β, could potentially be a kinase that phosphorylates STAT1-S727. This notion was supported by the physical association of mTOR with STAT1 in vivo, which was previously reported[29], although this association did not result in STAT1-S727 phosphorylation[29, 30]. We confirmed the co-IP of mTOR and STAT1 (Supplemental Fig. 3) and verified that this direct interaction indeed failed to phosphorylate STAT1-S727 also in our Karpas-299 cells, even though mTOR can phosphorylate STAT1 S727 in an *in vitro* kinase assay (Fig. 3b). We based this on the observation that STAT1-S727 phosphorylation was not reduced when rapamycin or AZD-8055 inhibited mTOR kinase activity (Fig. 3c). This result also excludes any downstream mTOR-activated kinases that could play a role in STAT1-S727 phosphorylation.

**Figure 3.**
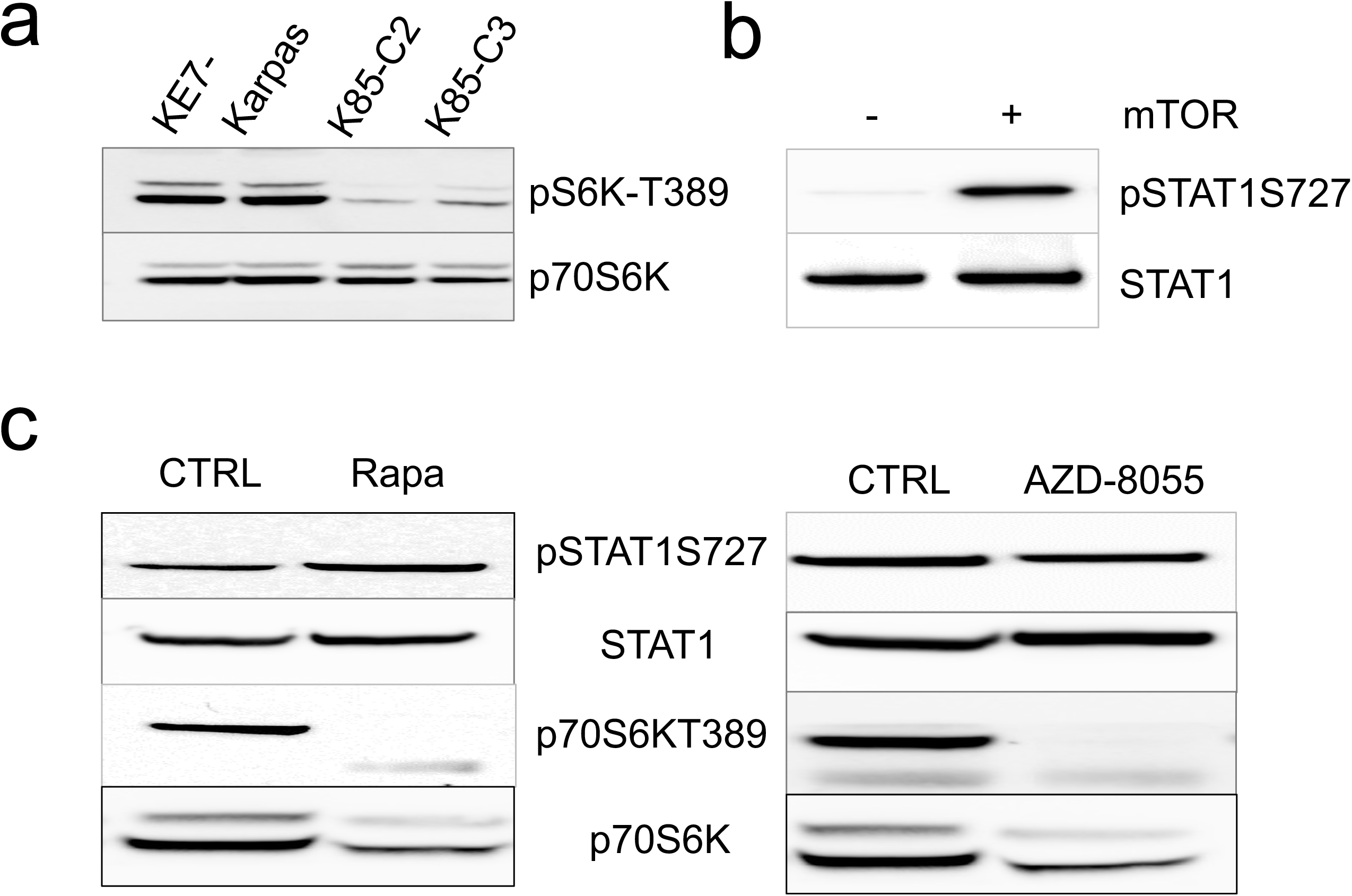
mTOR is not responsible for STAT1-S727 phosphorylation. (**a**) Immunoblot of lysates from equal numbers of Karpas-299 and K85 cells probed for pP70S6K-T389 and total P70S6K. (**b**) Immunoblot of an *in vitro* STAT1 kinase assay in the absence or presence of the mTOR kinase domain, probed for pSTAT1-S727. (**c**) Immunoblots of lysates from equal numbers of Karpas-299 cells, and Karpas-299 cells treated with rapamycin (1μg/ml for 3 days) (**left panel**) or AZD-8055 (100ng/ml for 3 days) (**right panel**), probed for pSTAT1-S727, STAT1, pP70S6K-T389, and total P70S6K.

### GSK3**β** is indirectly required for STAT1-S727 phosphorylation

Next, we investigated whether GSK3β is required for the phosphorylation of STAT1-S727. If GSK3β or a downstream kinase is required for the phosphorylation of STAT1-S727, the re-expression of GSK3β in K85 cells should restore STAT1-S727 phosphorylation. After transduction of a GSK3β lentiviral vector into K85 cells (called K85+GSK3), STAT1-S727 phosphorylation was restored (Fig. 4a), strengthening the link between GSK3β and STAT1-S727 phosphorylation. Furthermore, this also restored the expression of ETV7 (Fig. 4a), consistent with our previous conclusion that activated STAT1 is essential for the expression of ETV7. Furthermore, STAT1-S727 was partially inhibited by the GSK3β-specific inhibitor, Chir-98014 (Fig. 4b), which confirms that GSK3β is an upstream kinase responsible for STAT1-S727 phosphorylation in Karpas-299 cells. Partial inhibition of GSK3β was confirmed by the partial inhibition of pβ-Catenin-S33/37/T41 and resulted in reduced ETV7 protein and mRNA expression (Fig. 4b, 4c). If STAT1 is a direct physiological substrate of GSK3β, the phosphorylation of STAT1-S727 should change in parallel with the alteration in GSK3β activity. Therefore, we performed a GSK3β inhibitor time course using TDZD-8 in Karpas-299 cells and compared the inhibition of GSK3β activity with the level of STAT1-S727 phosphorylation at different time points. As shown in Figure 4d, GSK3β was already partially inhibited after two hours of TDZD-8 treatment and remained the same at 5 hours of treatment, as monitored by phosphorylation of pβ-catenin-S33/37/T41. However, the phosphorylation level of STAT1-S727 remained unaffected during this 5-hour time course, but was reduced after 18 hours of treatment. This suggested that a much longer time of inhibition of GSK3β results in reduced STAT1-S727 phosphorylation and that STAT1 is not a direct target of GSK3β kinase, even though GSK3β did phosphorylate STAT1-S727 in an *in vitro* kinase assay (Supplemental Fig. 4) and STAT1 and GSK3β physically interacted[31]. Overall, our results suggest that GSK3β is indirectly required for the phosphorylation of STAT1-S727, which in turn is essential for ETV7 expression, but it does not directly phosphorylate STAT1-S727.

**Figure 4.**
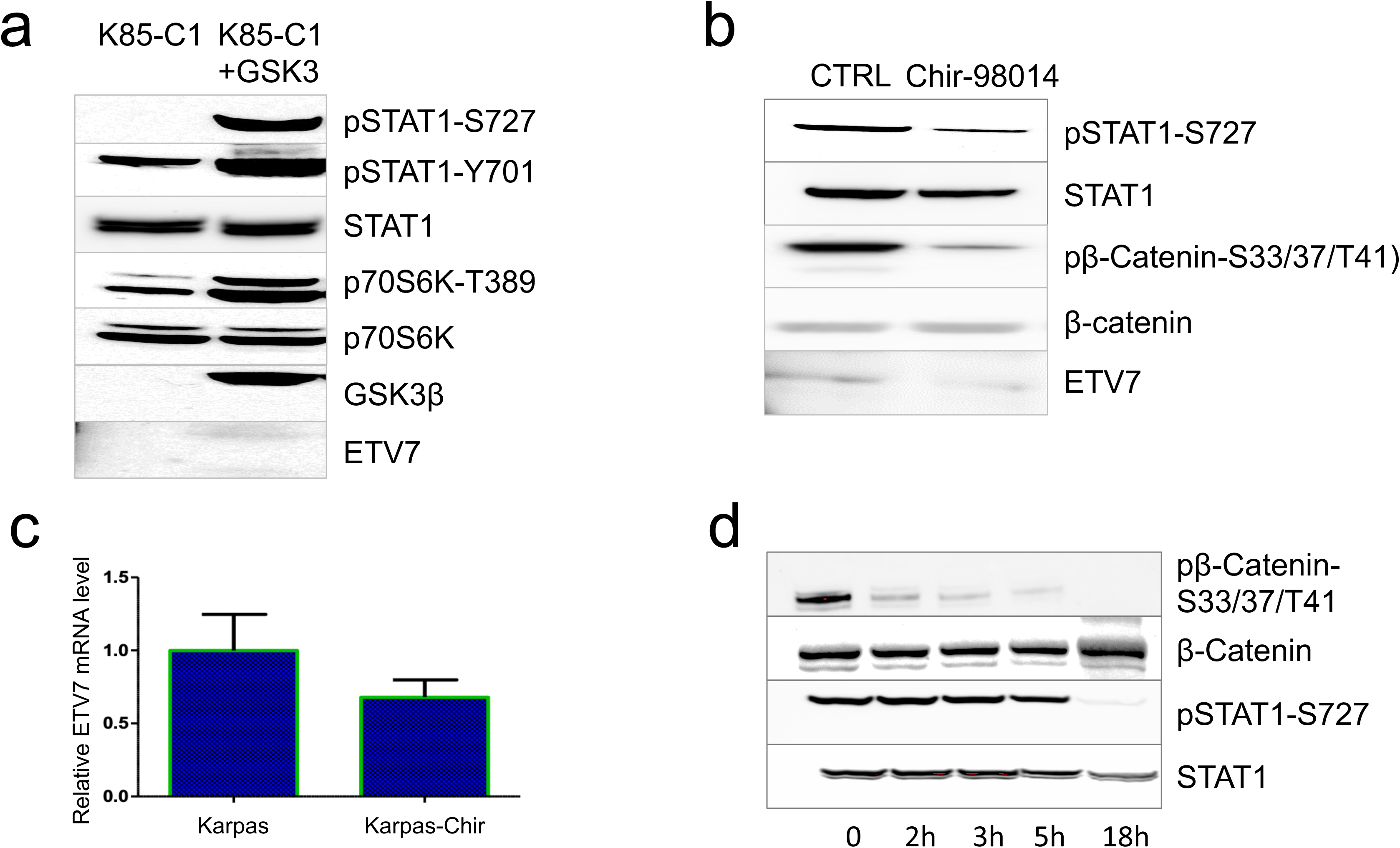
GSK3β is required for STAT1-S727 phosphorylation. (**a**) Immunoblots of lysates from equal numbers of K85-C1 and K85-C1+GSK3β cells probed for STAT1-S727, pSTAT1-Y701, p70S6K-T389, GSK3β, and ETV7. (**b**) Immunoblots of lysates from equal numbers of Karpas-299 and Karpas-299 cells treated with the GSK3β inhibitor Chir-90814 (2μM for 3 days), probed for pSTAT1-S727 and STAT1, pCTNB1-S33/37/T41) and CTNB1. (**c**) ETV7 mRNA levels in Karpas-299 and Karpas-299 cells treated by Chir-90814 (2μM for 3 days), as determined by real-time RT-PCR. (**d**) Karpas-299 cells were treated with the GSK3β inhibitor TDZD-8 (10μM) for 0, 2, 3, 5, and 18 hours. Immunoblots of lysates from equal numbers of cells were probed for pSTAT1-S727, STAT1, pCTNB1-S33/37/T41, and CTNB1.

### PKC**δ** is responsible for the phosphorylation of STAT1-S727

Downstream of GSK3β two kinases have been reported that phosphorylate STAT1-S727, PKCδ[32] [33] and SGK1[34]. While inhibiting SGK1, as monitored by SGK1-S422 phosphorylation, STAT1-S727 phosphorylation remained unaltered (Supplemental Fig. 5a), suggesting that SGK1 does not play a role in this phosphorylation event. Furthermore, SGK1 activity does not correlate with STAT1-S727 phosphorylation in Karpas-299 and K85-C1 cells (Fig. 5a). To investigate whether PKCδ contributes to STAT1-S727 phosphorylation, we treated Karpas-299 and K85-C1 cells with PMA, a PKC activator and found that pSTAT1-S727 increased significantly in Karpas-299, but not in K85-C1 cells (Fig 5a). Concurrently, PMA treatment of cell cultures increased pPKCδ-Y311 phosphorylation in Karpas-299 cells but not in K85-C1 cells, suggesting a link between PKCδ activation and STAT1-S727 phosphorylation. This also indicated that GSK3β is upstream of PKCδ and therefore required for its activation. As a control that PMA activation works in K85-C1 cells, we showed that there was an increase in ERK1/2 phosphorylation (Fig. 5a). Using the PKC inhibitor Go6983, we found that PMA-induced STAT1-S727 phosphorylation was blocked (Fig. 5b). The inhibition of PKCδ was confirmed by loss of PKCδ-Y311 phosphorylation (Fig. 5b), which further strengthened the potential role of PKCδ in STAT1-S727 phosphorylation in Karpas-299. As GSK3β is an upstream kinase of PKCδ, we also observed activation of GSK3β after PMA treatment, as indicated by increased phosphorylation of pβ-Catenin-S33/37/T41 and GSK3β-Y216 (supplemental Fig. 5b). Furthermore, PKCδ can phosphorylate STAT1-S727 *in vitro* (supplemental Fig. 5c). Finally, we followed changes in STAT1 and PKCδ phosphorylation during a time course of PMA treatment. Within 10 minutes STAT1-S727 and GSK3β-Y216 were already fully phosphorylated simultaneously (Fig. 5c). Taken together, we conclude that PKCδ is the kinase that phosphorylates STAT1-S727 in Karpas-299 cells *in vivo*, which is consistent with studies published by others[35, 36] using different cell lines.

**Figure 5.**
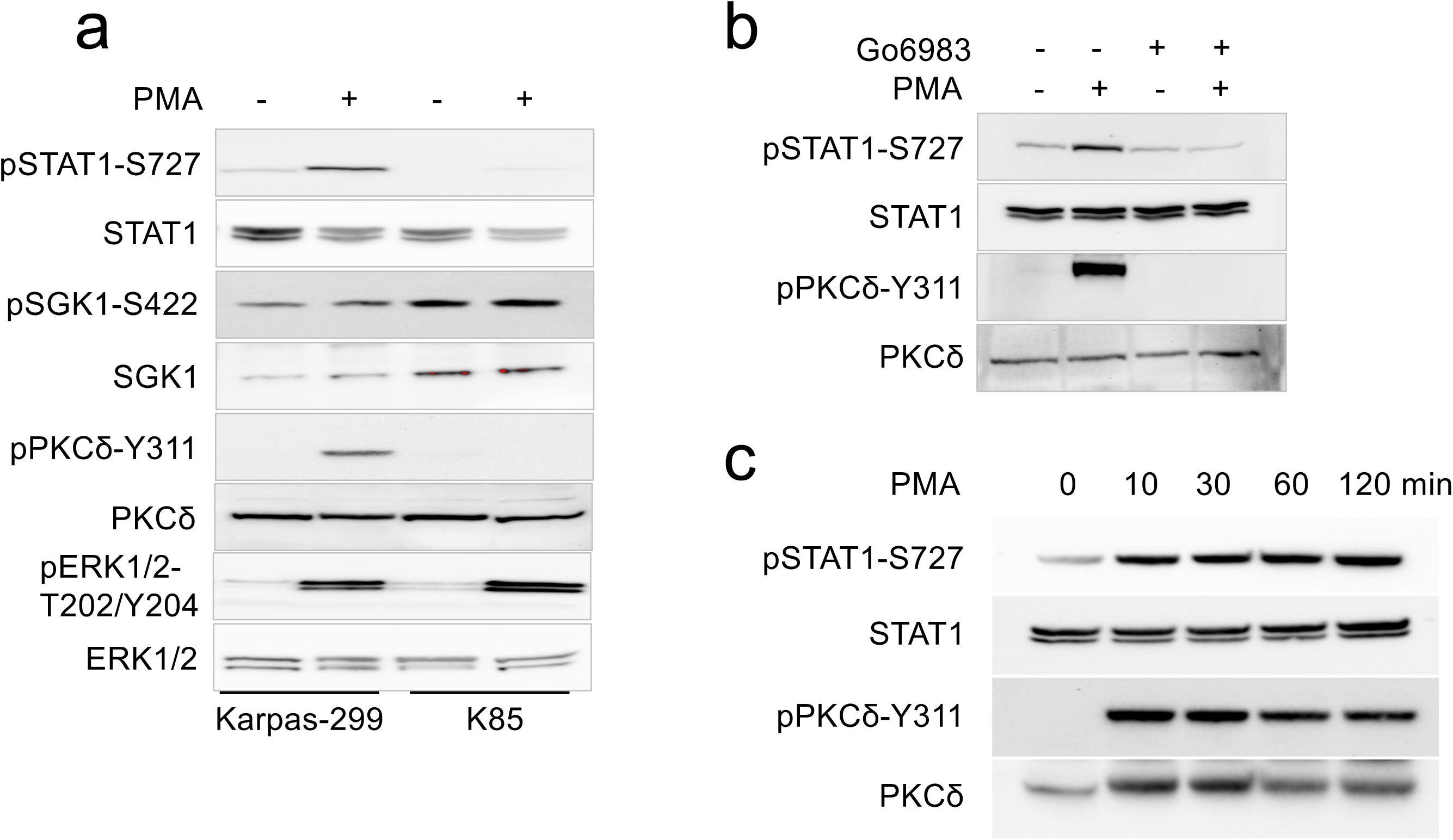
PKCδ regulates STAT1-S727 phosphorylation. (**a**) Immunoblot of lysates from equal numbers of Karpas-299, Karpas-299 treated with PMA (1μM) for 45minutes, K85-C1, and K85-C1 cells treated with PMA (1μM) for 45min, probed for pSTAT1-S727, STAT1, SGK1-S422, SGK1, PKCδ-Y311, PKCδ, ERK-T202/Y204, and ERK1/2. (**b**) Immunoblot of lysates from equal numbers of Karpas-299, Karpas-299 treated for 30 minutes with PMA (1μm), Karpas-299 treated for 1 hour with the PKC inhibitor Go6983 (10μm), Karpas-299 treated for 1 hour with the PKC inhibitor Go6983 (10μm) with the addition of PMA (1μm) for the last 30 minutes, probed for STAT1, pSTAT1-S727, PKC δ-Y311, PKCδ. (**c**) Immunoblot of lysates from equal numbers of Karpas-299 cells treated for 0, 10, 30, 60, and 120 minutes with PMA (1μm) was probed for pSTAT1-S727, STAT1, pPKCδ-Y311, and PKCδ.

## Discussion

We found that mTORC3 activity was affected by GSK3β, by disrupting GSK3β in Karpas-299 cells. Loss of GSK3β led to poor cell growth, in part due to loss of ETV7, an essential component of mTORC3, which stimulates cell growth. The poor cell growth is consistent with their slow cell cycle progression as indicated by a reduced percentage of K85 cell clones in S-phase compared with that of Karpas-299 cells but resembling that of the ETV7 deficient KE7-cells. This indicated that ETV7 indeed contributes to the slow cell growth, and that GSK3 plays an additional role in the cell growth. The deletion of GSK3β also abolished STAT1 activity, as indicated by the loss of its phosphorylation of S727, a post-translational modification essential for its transactivation activity. Through various experiments, we confirmed that STAT1 is a transcriptional activator of the *ETV7* gene. Therefore, GSK3β indirectly influences the activity of mTORC3 by regulating the transcriptional activity of STAT1.

The transcriptional activity of STAT1 is regulated by phosphorylation of both Y701 and S727. STAT1-Y701[37, 38] is phosphorylated by JAK kinases in the cytoplasm, leading to a reorientation of subunits within the STAT dimer, followed by translocation to the nucleus where it functions as a transcription factor[39, 40]. Early during activation, a second, independent phosphorylation event occurs at S727. The phosphorylation of STAT1-S727 is required for nearly 80% of its IFN-induced transcriptional activity[27, 28, 41]. Although numerous kinases have been claimed to phosphorylate STAT1-S727, such as CDK8[41], FAK[42], PKCδ[32], and SGK1[34], the kinase responsible for the phosphorylation of STAT1-S727 had not been truly resolved.

We show that GSK3β is an upstream kinase of STAT1-S727. Disruption of GSK3β in Karpas-229 cells abolished STAT1-S727 phosphorylation, while re-expression of GSK3β restored this activity. In addition, GSK3β inhibition reduced STAT1-S727 phosphorylation, further indicating that GSK3β activity is required for this modification. This is consistent with previous work published by others; Firstly, GSK3β was shown to be activated by IFN-γ signaling[43], and activated GSK3β was shown to synergistically facilitate IFNγ-induced STAT1 activation[43]. Furthermore, GSK3β knockdown blocked the IFN-α-induced phosphorylation of STAT1[44]. In addition, GSK3β inhibitors were found to inhibit Th1 T cell production by reducing STAT1 activation [45].

Together, this indicated the importance of GSK3β activity for STAT1 activation, but direct phosphorylation of STAT1 by GSK3β had never been reported. To establish that a potential substrate is a true physiological substrate of GSK3β is not straightforward; however, a minimum requirement is that the substrate phosphorylation (STAT1-S727) changes in parallel with changes in GSK3β activity. We did not see a change in STAT1-S727 phosphorylation immediately after inhibition of GSK3β activity, suggesting that STAT1-S727 is not a direct substrate of GSK3β. Therefore, it was likely that a kinase downstream of GSK3β directly phosphorylates STAT-S727.

It has been reported that PKCδ is a kinase downstream of GSK3β[46]. PKCδ is a member of the PKC family of serine-threonine kinases, which play important roles in signaling downstream of various cytokine receptors [47, 48]. Therefore, we investigated whether instead of GSK3β, its downstream kinase PKCδ phosphorylates STAT1-S727. Like GSK3β, PKCδ can be activated by IFN-α or INF-γ[33, 49], and as a result, it can phosphorylate STAT1-S727[33]. Besides using specific pharmacological inhibitors of PKCδ, or a dominant-negative PKCδ mutant, luciferase reporter transcription assays were also used to support this conclusion[33]. Unlike GSK3β, PKCδ only affects STAT1-S727 but not STAT1-Y701 phosphorylation.

Many studies have reported that INF signaling activates GSK3β or PKCδ resulting in STAT1 phosphorylation and activation. However, in Karpas-299 cells IFNγ has only a minor effect on STAT1 phosphorylation. In Karpas-299 cells, PMA activates PKCδ and in turn, phosphorylates STAT1-S727, but we did not observe an increase in the expression of ETV7. We believe this is because STAT1 activity requires dual STAT1-S727 and STAT1-Y701 phosphorylation, and PKCδ targets STAT1-S727 only. As PKCδ activity correlated with the STAT1-S727 phosphorylation in untreated and PMA-treated cells, we believe that both basic level STAT1-S727 phosphorylation and PMA-activated STAT1-S727 phosphorylation are mediated by PKCδ.

Together, our work provides evidence that GSK3β controls mTORC3 activity. We identified a novel pathway in which GSK3β, through PKCδ/STAT1, regulates ETV7 transcription and thus positively regulates the activity of mTORC3. The identification of this GSK3β/PKCδ/STAT1/ETV7/mTORC3 signaling pathway could benefit the design of therapeutics for numerous diseases, including cancer.

## Acknowledgments

We thank Diantha van de Vlekkert for help with the STAT1 C&R/Q-PCR experiment. Support for this work was provided by the Cancer Center Support Grant CA 021765 and the American Lebanese Syrian Associated Charities (ALSAC) of St Jude Children’s Research Hospital.

## Authorship

Contribution: J.Z designed and performed the research and wrote the manuscript; F.C.H generated the K85 and STAT1 cell clones as well as Karpas-299 cell clones; M.C. performed the C&R/Q-PCR experiments; G.C.G developed the research plan, supervised the research, and edited the manuscript.

## Data availability statement

Additional data are available upon request from the authors.

Correspondence to Gerard C Grosveld and Jun Zhan

