## Supplemental data for "mTORC3 activity is regulated by GSK3β through the activation of PKCδ that phosphorylates and activates STAT1"

Supplemental Materials:

Supplemental Figures:

a.

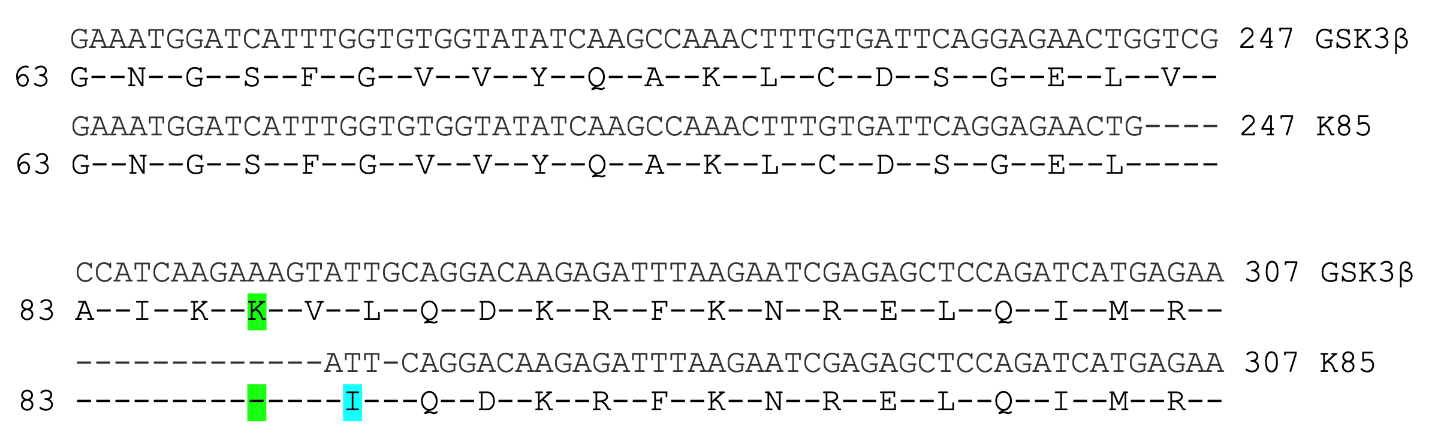

b.

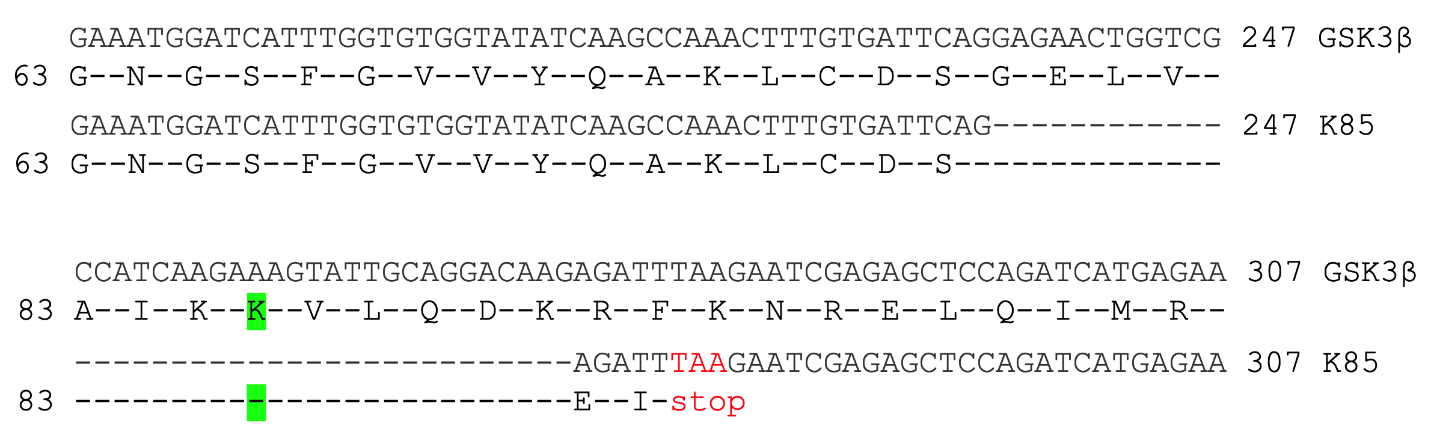

**c.**

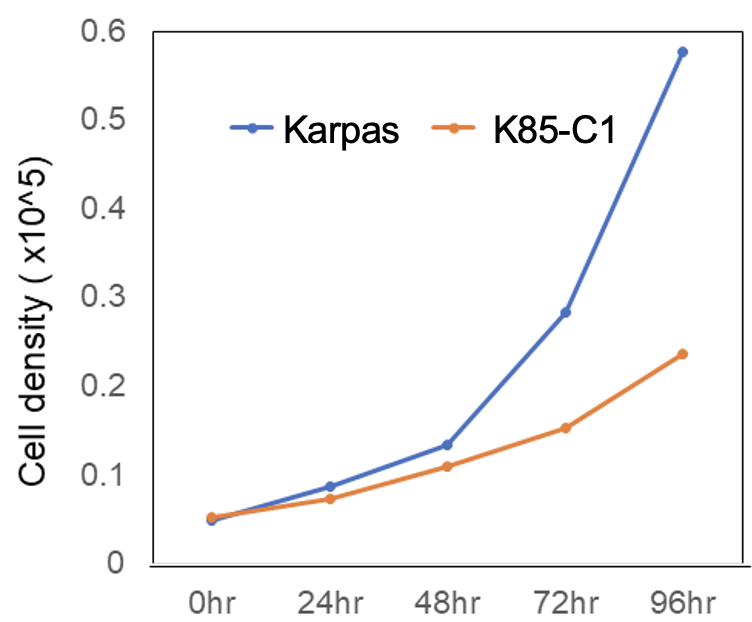

**d.**

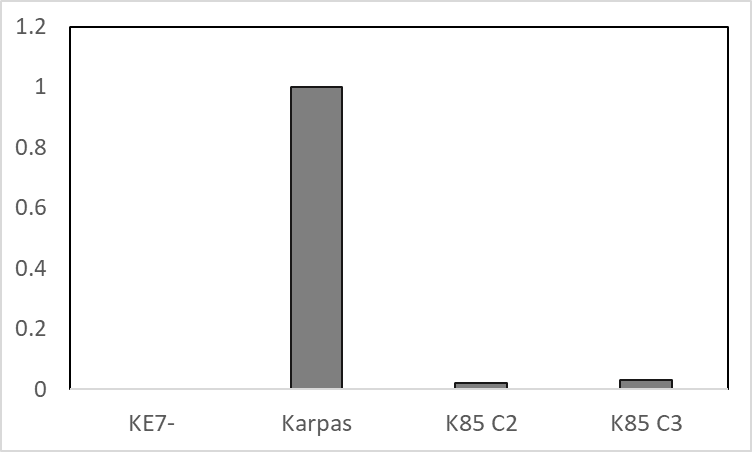

Relative ETV7 mRNA level

|  |  |
| --- | --- |
| KE7- | 0 |
| Karpas | 1 |
| K85 C2 | 0.02 |
| K85 C3 | 0.03 |

**e**

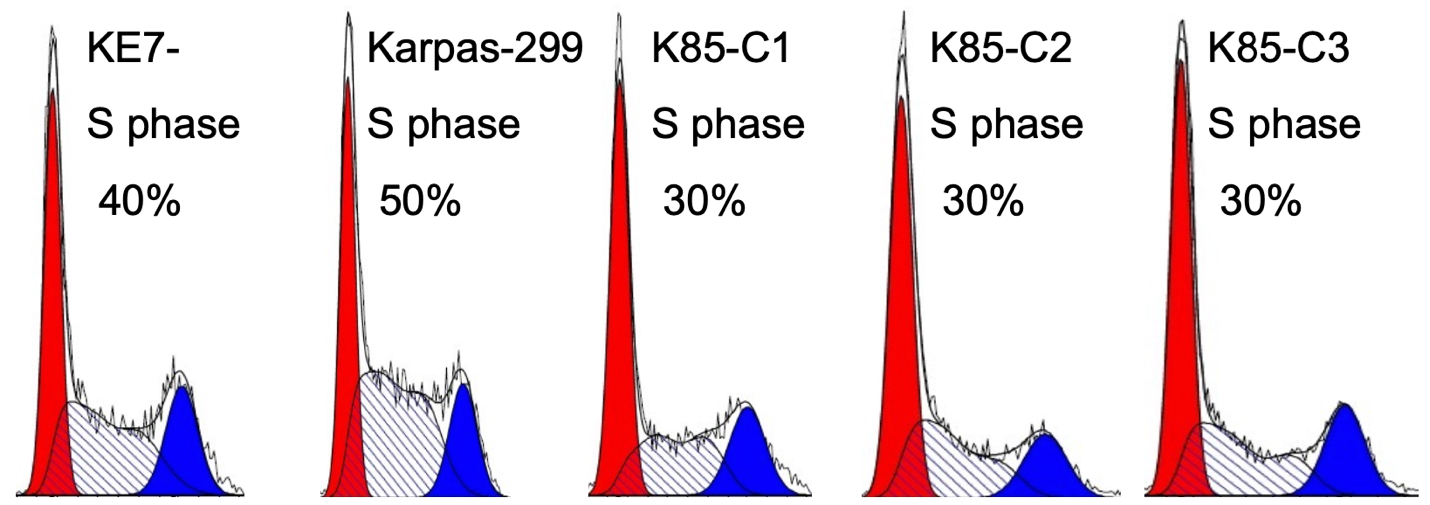

Supplemental Figure 1.

a. cDNA nucleotide and amino acid sequence of the region covering the Cas9 targeted region of GSK3β in the K85-C1 cell line. Depicts a 17 bp deletion around K85 in K85 cells and an additional single bp deletion 4 bp downstream, leading to an in-frame deletion of 6 amino acids including K85, and an insertion of 1 nonhomologous amino acid (isoleucine). K85-C1 is highlighted in green, and the nonhomologous isoleucine is highlighted in blue. The depicted GSK3β cDNA sequence shown starts at bp 187/amino acid 63.

b. A 38 bp deletion around K85 leading to a frameshift and a stop 2 codons later. The stop codon is indicated in red.

c. Growth curve of K85-C1 compared with its parent line Karpas-299.

d. Relative amounts of ETV7 mRNA in Karpas-299, KE7- and two K85 cell clones (C2 and C3) were determined by real-time RT-PCR

e. Flow cytometric cell cycle analysis of KE7-, Karpas-299, and different K85 cell clones (K85-C1, K85-C2 and K85-C3)

a.

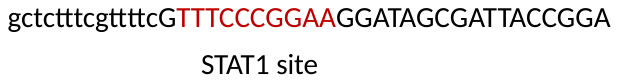

b

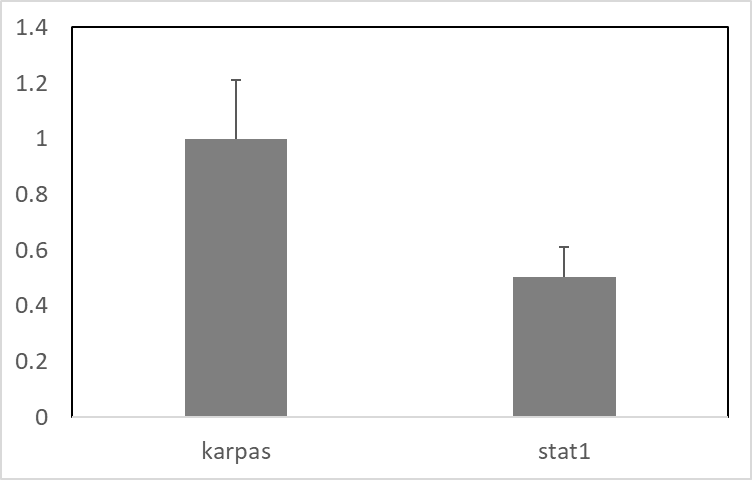

Relative ETV7 mRNA level

|  |  |
| --- | --- |
| karpas | 1 |
| stat1 | 0.503802 |

c.

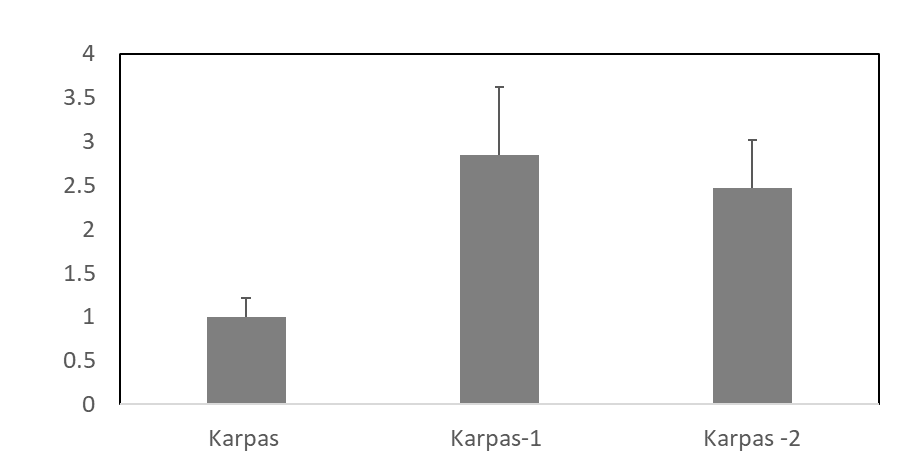

Relative ETV7 mRNA level

|  |  |
| --- | --- |
| Karpas | 1 |
| Karpas-1 | 2.84194 |
| Karpas -2 | 2.46634 |

d.
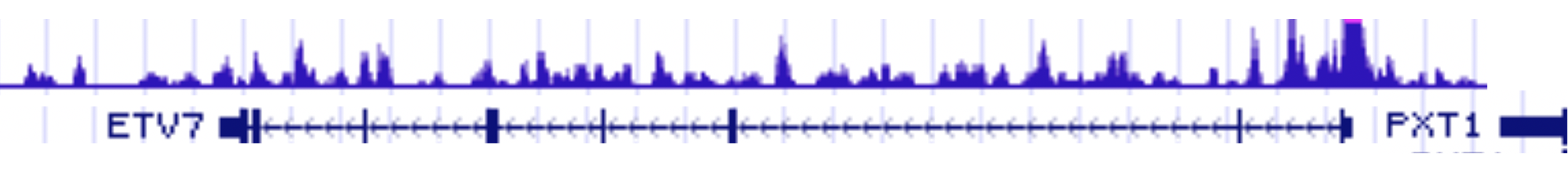

e.

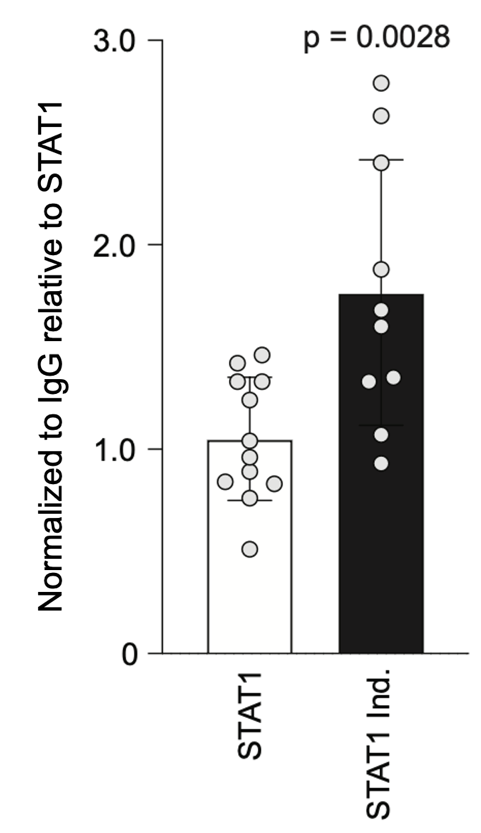

f.

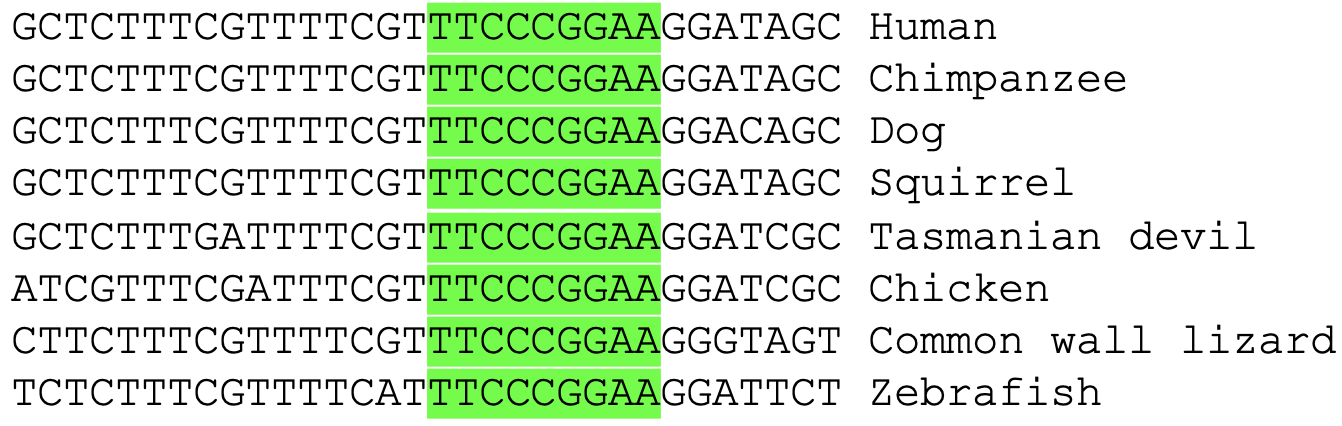

Supplemental Figure 2. STAT1 binds to the P2 ETV7 promoter.

a. Map of the ETV7 promoter area showing two promoters: P2 at position 0 and P1 at position +116 ([https://epd.epfl.ch//index.php](https://epd.epfl.ch/index.php)). Vertical bars depict the relative frequency of ETV7 transcriptional start sites (TSS) in different human tissues and cell lines. The two red bars represent the TSS at P2 (position 1) and P1 (position 116). The sequence of the STAT1 binding site at +6 bp is shown in red below the map.

b. Relative amounts of ETV7 mRNA in Karpas-299 and STAT1 mutated cells were determined by real-time RT-PCR.

c. Relative amounts of ETV7 mRNA in Karpas-299 and two different K85 clones (C2 and C3) were determined by real-time RT-PCR

d. STAT1 ChIP-seq analysis of the ETV7 gene in INFγ induced K562 cells as reported in ENCODE.

e. STAT1 Cut & Run/Q-PCR assay of the ETV7 promoter area of Karpas-299 cells induced or not for 30 minutes with INFγ. The data are the average of 4 independent experiments normalized against IgG.

f. P2 promoter sequences of different classes of vertebrates showing the fully conserved STAT1 binding site highlighted in green.

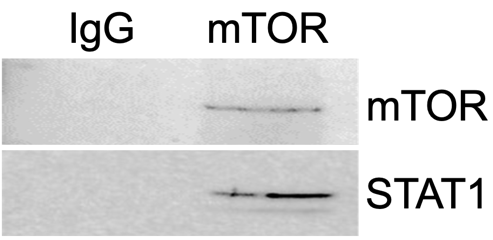

Supplemental Figure 3. The association of mTOR and STAT1 in vivo.

Immunoblot of Karpas-299 lysate immunoprecipitated with IgG or mTOR antibody probed for mTOR and STAT1.

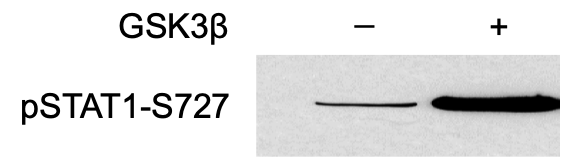

Supplemental Figure 4. *In vitro* kinase assay of STAT1 phosphorylated by GSK3β.

Immunoblot of an *in vitro* STAT1 kinase assay in the presence or not of active GSK3β, probed for pSTAT1-S727.

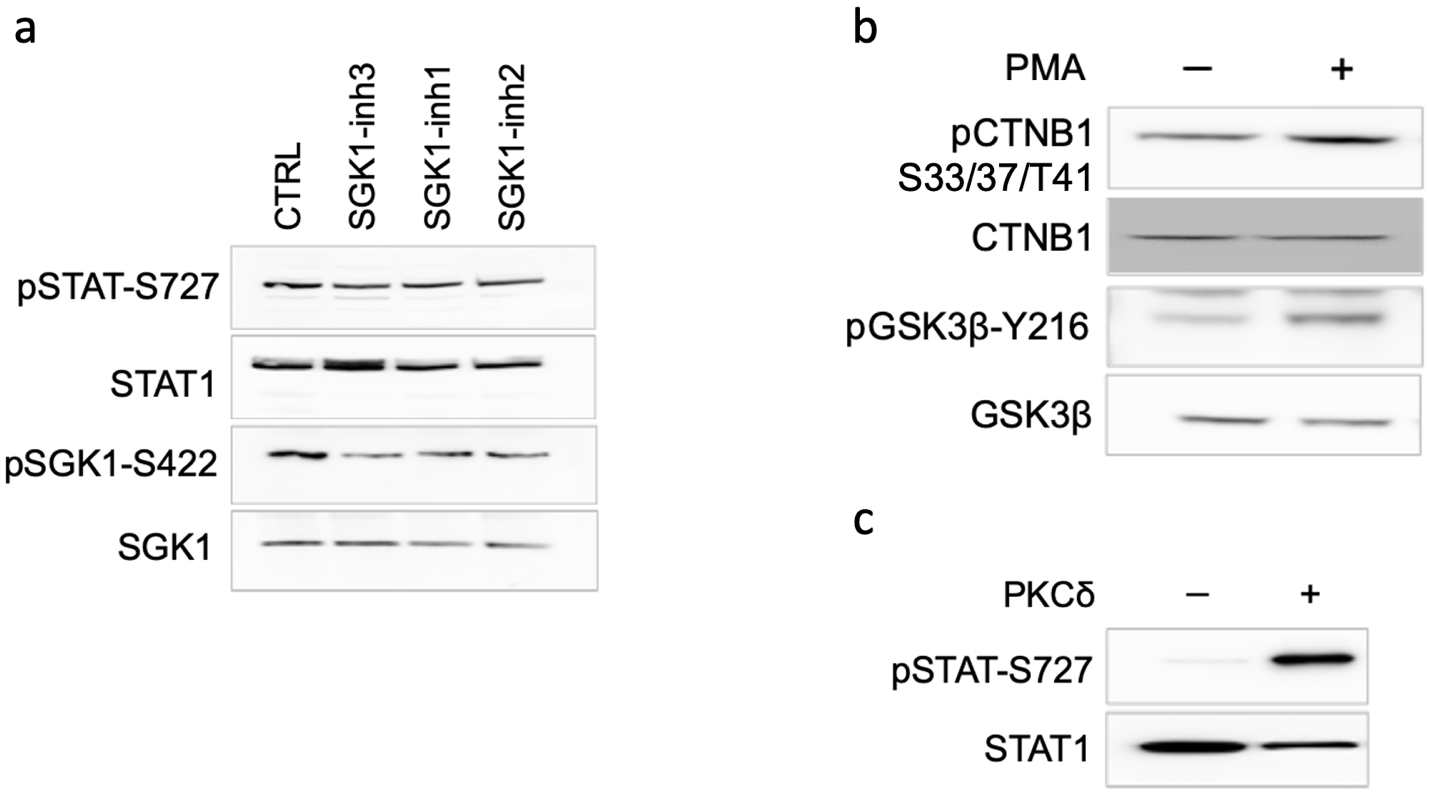

Supplemental Figure 5.

a. Immunoblots of lysates of equal numbers of Karpas-299 cells treated for 3 days with the SGK1 inhibitors SGK1-in-1(10μM), SGK1-in-2(10μM), and SGK1-in-3 (GSK650394) (10μM), probed for pSTAT1-S727, STAT1, pSGK1-S422, and SGK1.

b. Immunoblot of lysates of equal numbers of Karpas-299 cells treated for 30 minutes with PMA (1μm) and probed for pβ-Catenin-S33/37/T41, β-catenin, GSK3β, and pGSK3β-Y216.

c. Immunoblot of an *in vitro* kinase assay of STAT1 phosphorylated by active PKCδ and probed for pSTAT1-S727.
